# A high-throughput 384-well CometChip platform reveals a role for 3-methyladenine in the cellular response to etoposide-induced DNA damage

**DOI:** 10.1101/2022.03.29.486126

**Authors:** Jianfeng Li, Alison Beiser, Nupur B. Dey, Shunichi Takeda, Liton Kumar Saha, Kouji Hirota, L. Lynette Parker, Mariah Carter, Martha I. Arrieta, Robert W. Sobol

**Author notes:** **To whom correspondence should be addressed:** Robert W. Sobol, Ph.D. Department of Pharmacology, College of Medicine Mitchell Cancer Institute University of South Alabama 1660 Springhill Avenue, AL 36604.

## Abstract

The Comet or single-cell gel electrophoresis assay is the gold standard for analysis of cellular, nuclear genome damage. However, low throughput limits its application for large-scale studies. To overcome these limitations, a 96-well CometChip platform was recently developed that increases throughput and reduces variation due to simultaneous processing and automated analysis of 96 samples. To advance throughput further, we developed a 384-well CometChip platform that allows analysis of ∼100 cells per well. The 384-well CometChip extends the capacity by 4-fold as compared to the 96-well system, enhancing application for larger DNA damage analysis studies. The overall sensitivity of the 384-well CometChip is consistent with that of the 96-well system, sensitive to genotoxin exposure and to loss of DNA repair capacity. We then applied the 384-well platform to screen a library of protein kinase inhibitors to probe each as enhancers of etoposide induced DNA damage and found that 3-methyladenine significantly increased levels of etoposide-induced DNA damage. Our results suggest that a 384-well CometChip is useful for large-scale DNA damage analyses, which may have increased potential in the evaluation of chemotherapy efficacy, compound library screens, population-based analyses of genome damage and evaluating the impact of environmental genotoxins on genome integrity.

## INTRODUCTION

The Comet assay is the gold standard for the analysis of nuclear genomic DNA damage at the single cell level (1-3). The assay was originally developed by Ostling and Johanson for the analysis of DNA double-strand breaks under neutral conditions (4). It was then further developed by Singh et al. for the detection of alkali-labile sites and DNA single-strand breaks under highly alkaline conditions (pH>13) with high sensitivity (50–15,000 breaks/cell) (5). In a traditional alkaline comet assay, cells are embedded in agarose on a slide and the embedded cells are lysed and then the DNA is denatured. The slides are then subject to electrophoresis whereby the fragmented DNA rapidly migrates from the nucleoid body containing the intact DNA. Once the DNA is stained, the images reveal a distinct tail and head shape of the DNA, referred to as comets. The ratio of the DNA content in the tail, as compared to the content in the head, is proportional to the extent of DNA damage (3). The comet assay can be applied to a wide variety of eukaryotic cells from sources such as animal organs or plants as well as prokaryotic cells in either a cycling or noncycling phase, as well as in fresh or frozen status (6). The traditional Comet assay is a straightforward, relatively inexpensive, and highly sensitive method. However, the approach contains two major limitations for utilization in large-scale DNA damage analysis studies. The first is low throughput - in the traditional assay, only 20 slides containing one or two gels can be run in a single electrophoresis step (7). The second limitation is high variability for inter- or intra-laboratory experiments. This lack of reproducibility can be caused by the numerous variations among cell growth, treatment, lysis, denaturation, electrophoresis, imaging, and analysis conditions. The inter-sample variations can range from 10% to 20% (8) and the variations across different laboratories can vary up to 80% (9).

To overcome these limitations, a single-cell trapping microwell approach was developed (10) that formed the basis for the development of the 96-well CometChip assay (11). The single-cell trapping microwell and CometChip systems comprise a layer of agarose gel bound to GelBond film (10) or chemically bound to a glass slide or chip (11), on which an array of microwells of precise diameters (most commonly 30 μm) are generated from a microfabrication-based silicon mold (12). The 96 wells per chip of the 96-well CometChip platform significantly increased the throughput compared to the 2 wells per slide approach in the traditional comet assay (11). Also, due to the single-cell trapping approach used in the system (10), the 96-well CometChip provides uniform cell distribution in a single focal plane in the micro-patterned agarose array (11). Such an approach avoids the variability that can be seen with cells on different focal planes in the traditional comet slide system. The variation from imaging is also drastically reduced (10). The 96-well CometChip platform has been successfully applied for the evaluation of DNA damage (11) including the measurement of DNA double-strand breaks with or without DNA repair inhibitors (13) and H_2_O_2_ or IR induced DNA damage (14). A modified version of the CometChip, the EpiComet-Chip, has also been developed for the specific assessment of DNA methylation status (15). We have applied the 96-well CometChip platform to the analysis of a 74-compound library from the National Toxicology Program to evaluate induced DNA damage upon exposure of human cells (11). We also evaluated DNA damage in field-collected blood samples from turtles (6). Recently, a 96-well HepaCometChip was developed for screening DNA damage induced by bulky procarcinogens using immortalized HepaRG™ cells (16). Further, the recent use of CometChip for the comparison of human DNA repair kinetics after H_2_O_2_-induced oxidative damage demonstrates the CometChip platform’s utility in future epidemiological and clinical studies (17). With the drastic decrease in inter-sample variation in a 96-well CometChip assay, only 20 comets were necessary for a statistically significant detection of DNA damage induced by a DNA damaging agent for doses within the linear range (14). This is far fewer than the 100-comet standard for a traditional comet assay (14), further supporting the benefit of increasing the throughput of the CometChip assay.

To that end, we report here a 384-well CometChip platform that includes a 384-well plate “macrowell former” to generate 384 individual wells on the glass-based CometChip with an average of 100 microwells per well. The 384-well CometChip platform extends the capability of the CometChip assay 4-fold as compared to the 96-well CometChip system. The overall sensitivity of the 384-well CometChip assay is like that of the 96-well CometChip assay (11). We demonstrate that the 384-well CometChip system clearly detected a dose-responsive increase in DNA damage caused by several genotoxins and showed the predicted enhancement of DNA damage following the loss (knockout) of the DNA repair scaffold gene X-ray repair cross complementing 1 (XRCC1). To demonstrate the applicability of a 384-well CometChip for large-scale DNA damage analysis, we used the system to screen a library of protein kinase inhibitors, evaluating potential enhancement of etoposide induced damage. Our findings indicate 3-methyladenine (3-MA) increased the level of DNA damage resulting from etoposide treatment. Overall, our results demonstrate that the 384-well CometChip is useful for large-scale DNA damage studies, which may have increased potential in the evaluation of chemotherapy efficacy, compound library screens, population-based analyses of genome damage and in the evaluation of the impact of environmental genotoxins on genome integrity.

## MATERIALS AND METHODS

### Compounds

Methyl methane sulfonate (MMS) and etoposide were obtained from Sigma-Aldrich (St. Louis, MO). The kinase inhibitor compound set was purchased from Cayman Chemical (Cat# 10505). All compounds were stored at −20°C until used. The final concentration of each kinase inhibitor used in the analysis was 10 μM.

### Cell Culture

TK6 cells were purchased from ATCC (ATCC® CRL-8015™). The TK6/XRCC1-KO cell line was generated by a pair of TALEN expression plasmids targeted to the XRCC1 gene as in the previous report (18). All the cells were cultured in RPMI-1640 media with 10% fetal bovine serum, supplemented with 1% penicillin-streptomycin at 5% CO_2_ at 37°C.

### CometChip System and Supplies

The 96-well CometChip system, described previously (11), was purchased from Bio-Techne (Cat#: 4260-096-CSK) including disposable 30 μm CometChips (Cat#: 4260-096-01), the CometChip 96-well magnetically sealed cassettes (formers), the CometChip Electrophoresis System (Cat# 4260-096-ESK), and the Comet Analysis Software (CAS). Blank CometChips needed for the 384-well system, with no painted wells, were purchased from Bio-Techne as a custom order.

### 384-well CometChip Former

To accommodate the 384-well approach, the 96-well former of the upper portion of a CometChip 96-well former cassette (donated by Trevigen), was hollowed out and was replaced with a bottomless 384-well plate (Greiner Bio-One, Cat# 781000) and epoxy sealed (**Figure 1**). Combined with the bottom portion of the 96-well magnetically sealed cassette (the former), the new 384-well former was then combined with a non-painted ‘blank’ CometChip (Bio-Techne) for the assay.

**Figure 1.**
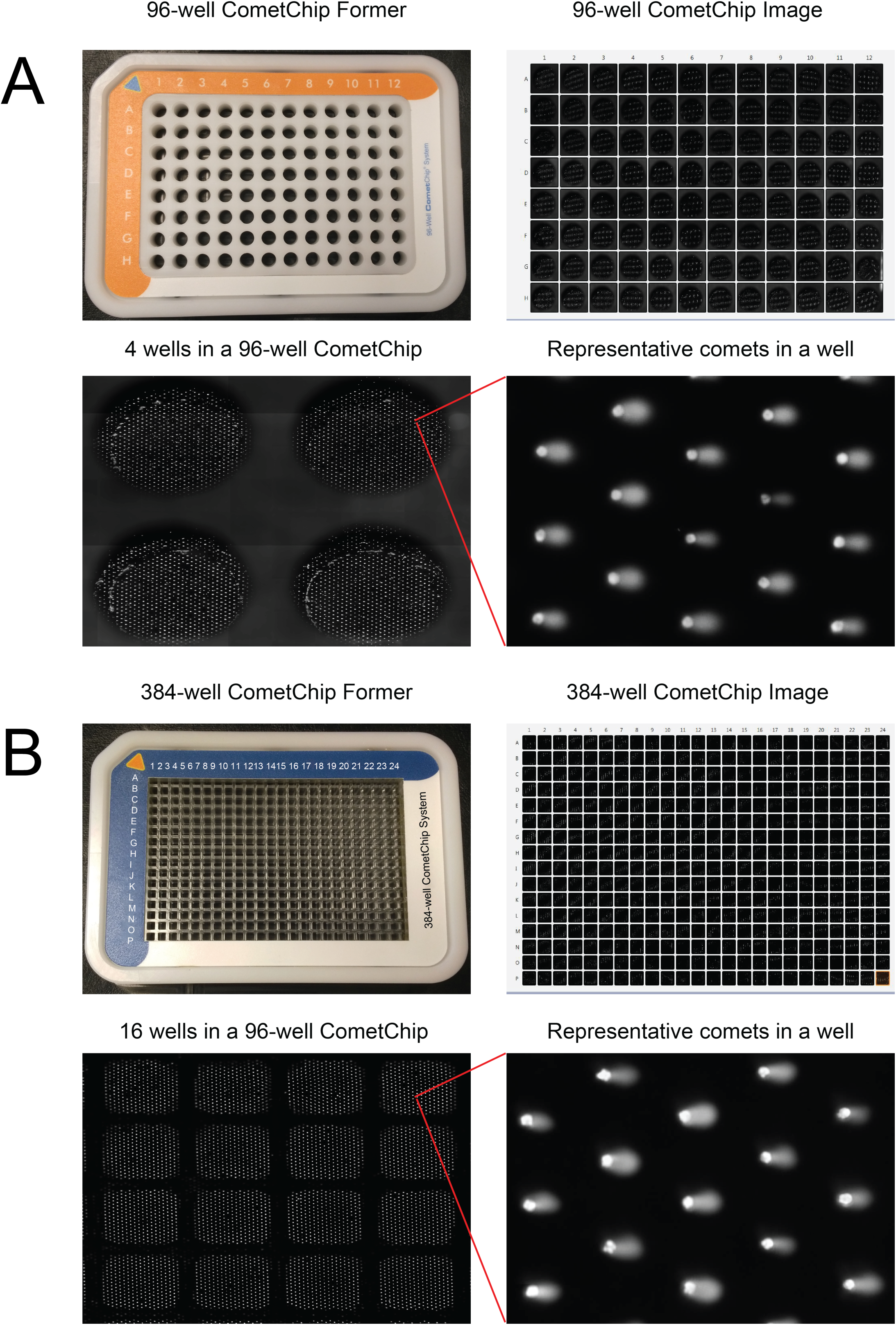
Development of the 384-well CometChip. (**A**) Top left: A 96-well CometChip former (Bio-Techne); Top right: an outline image of the entire 96-well CometChip; Bottom left: an image of comets in 4 wells of a 96-well CometChip; Bottom right: images of comets within one well of a 96-well CometChip. (**B**) Top left: A 384-well CometChip former; Top right: an outline image of the entire 384-well CometChip; Bottom left: an image of comets in 16 wells of a 384-well CometChip; Bottom right: images of comets within one well of a 384-well CometChip.

### CometChip Adapter for the Integra VIAFLO384 Electronic Pipette System

To accommodate the use of an electronic pipette system, we developed an adapter for the CometChip former (**Figure S1**). A plexiglass plate was designed to mount the CometChip former to the Integra VIAFLO384 Electronic Pipette, allowing for the precise orientation for the pipettor.

### CometChip Assay

The CometChip assay was performed as we described previously with minor modifications (11). TK6 or TK6/XRCC1-KO cells were seeded into a 96-well V-bottom microplate (100,000 cells/well) in growth medium (100 μl). After 1 hour of incubation, 100 μl of growth medium containing a 2x concentration of the DNA damaging agent (MMS or etoposide) was added into each well and the dish was incubated for 30 minutes at 37°C in the cell culture incubator. Subsequently, the cells were transferred to the corresponding well of the assembled CometChip apparatus. The cells settled into the microwells by gravity by placing the loaded CometChip apparatus for 20 min at 4°C. Then, the CometChip was washed with PBS to remove cells that had not settled into the microwells. Next, the CometChip was sealed with low melting point agarose (LMPA) (Topvision; ThermoFisher Scientific) (7 ml; 0.8% LMPA/PBS). The CometChip was then submerged in lysis solution with detergent (Bio-Techne) overnight at 4°C. After equilibration twice in 200 ml alkaline running buffer (200 mM NaOH, 1 mM EDTA, 0.1% Triton X-100, pH>13), the CometChip was electrophoresed at 22 V for 50 min at 4°C. This was followed by a re-equilibration to neutral pH twice in 100 ml Tris buffer (0.4 M Tris·Cl, pH 7.4) and twice in 100 ml Tris buffer (0.02 M Tris·Cl, pH 7.4) for 15 minutes each. Then, the DNA was stained with 1× SYBR Gold (ThermoFisher Scientific), diluted in Tris buffer (20 mM Tris·Cl, pH 7.4) for 30 min and de-stained twice in Tris buffer (20 mM Tris·Cl, pH 7.4) for 15 minutes.

### Automated Image Acquisition

The entire 96-well or 384-well CometChip was automatically imaged using the Celigo S imaging cytometer (Nexcelom Bioscience, Lawrence, MA) at the same image acquisition setting to avoid imaging variability. One image was generated for each well of the 96-well or 384-well CometChip at a resolution of 1 micron/pixel. The DNA damage (% Tail DNA) was automatically quantified from the comet images using the CometChip Analysis Software (Bio-Techne) with the 4x setting. Next, the quantitative data was exported to Excel (Microsoft) and subsequently to Prism 9 (GraphPad Prism) for plotting and statistical analysis.

### Kinase Inhibitor Screen

TK6 cells were seeded into four 96-well V-bottom microplates at a concentration of 100,000 cells per well in 100 μl growth medium and incubated for 1 hour. The inhibitors (**Table 1**), in two 96-well microplates of the kinase inhibitor library (Cayman Chemical, cat# 10505), were first diluted into each well of two new 96-well microplates to a 4x concentration (40 μM) using the TK6 cell growth medium just before treatment. Then 50 μl of the diluted kinase inhibitors (40 μM) from each of the two inhibitor plates were transferred into each well of two cell-containing microplates using the Integra VIAFLO384 Electronic Pipette system (INTEGRA Biosciences Corp, Hudson, NH 03051) as one group. The final concentration of each inhibitor after dilution into the cell containing wells was 10 μM. The procedure was repeated once so that the four cell-containing plates with the addition of kinase inhibitors were divided into two groups for the following experiment: one for the etoposide treatment group and one the control group with only the kinase inhibitors. Each group has two microplates matching the inhibitor plates. The plates were kept for 30 minutes at 37°C. Subsequently, etoposide containing growth medium (50 μl, 8 μM) was added into each well of the two microplates in the etoposide treatment groups. Additionally, DMSO containing growth medium (50 μl) was added into each well of two microplates in the control groups. All the plates were incubated for an additional 30 minutes at 37°C. The cells from each 96-well microplate were then transferred into each quarter of the assembled 384-well CometChip apparatus. Cells settled into the microwells by gravity after placing the loaded CometChip apparatus for 20 min at 4°C. The subsequent steps are the same as the 96-well CometChip assay for lysing, electrophoresing, imaging, and quantifying the comets.

**TABLE 1.**
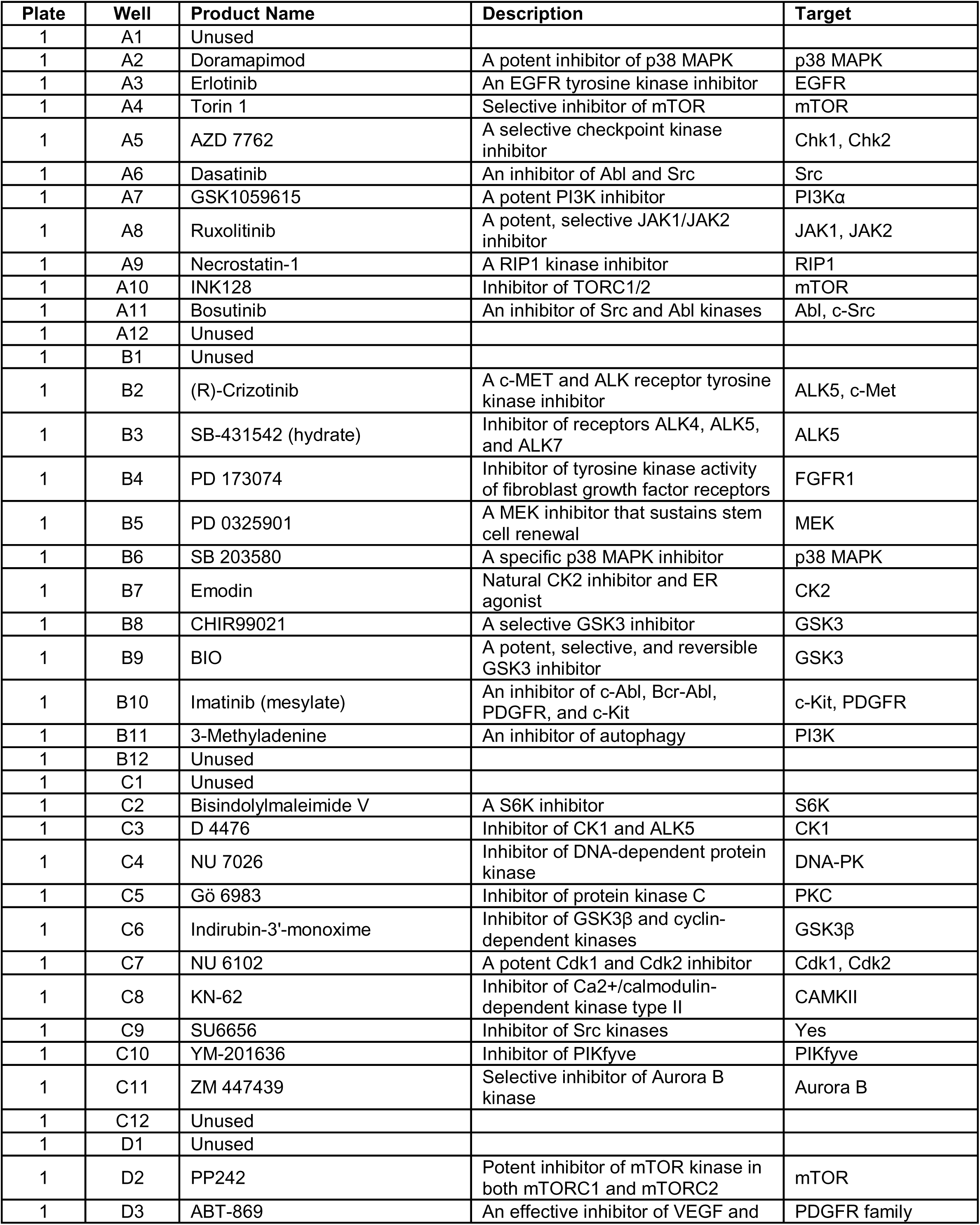
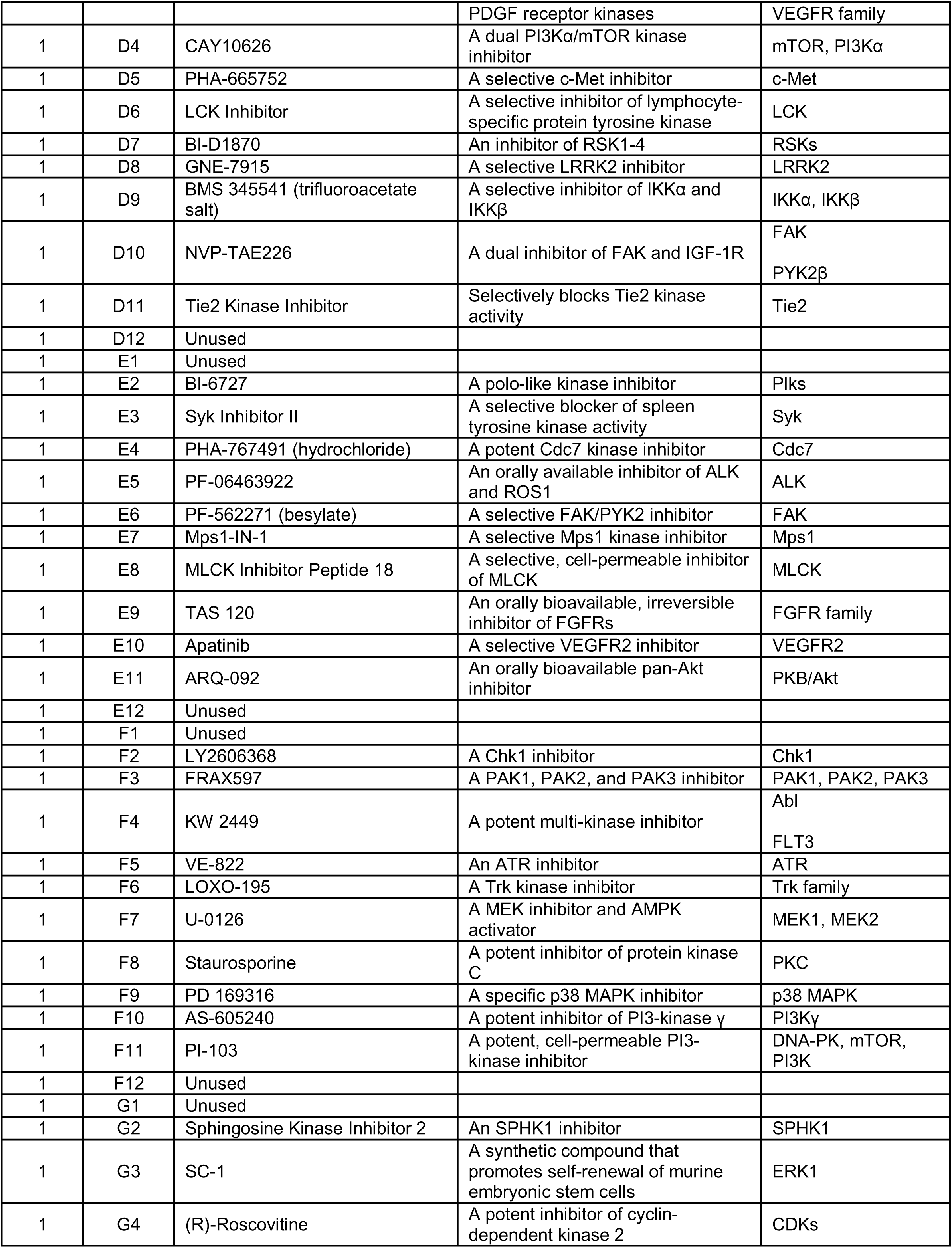
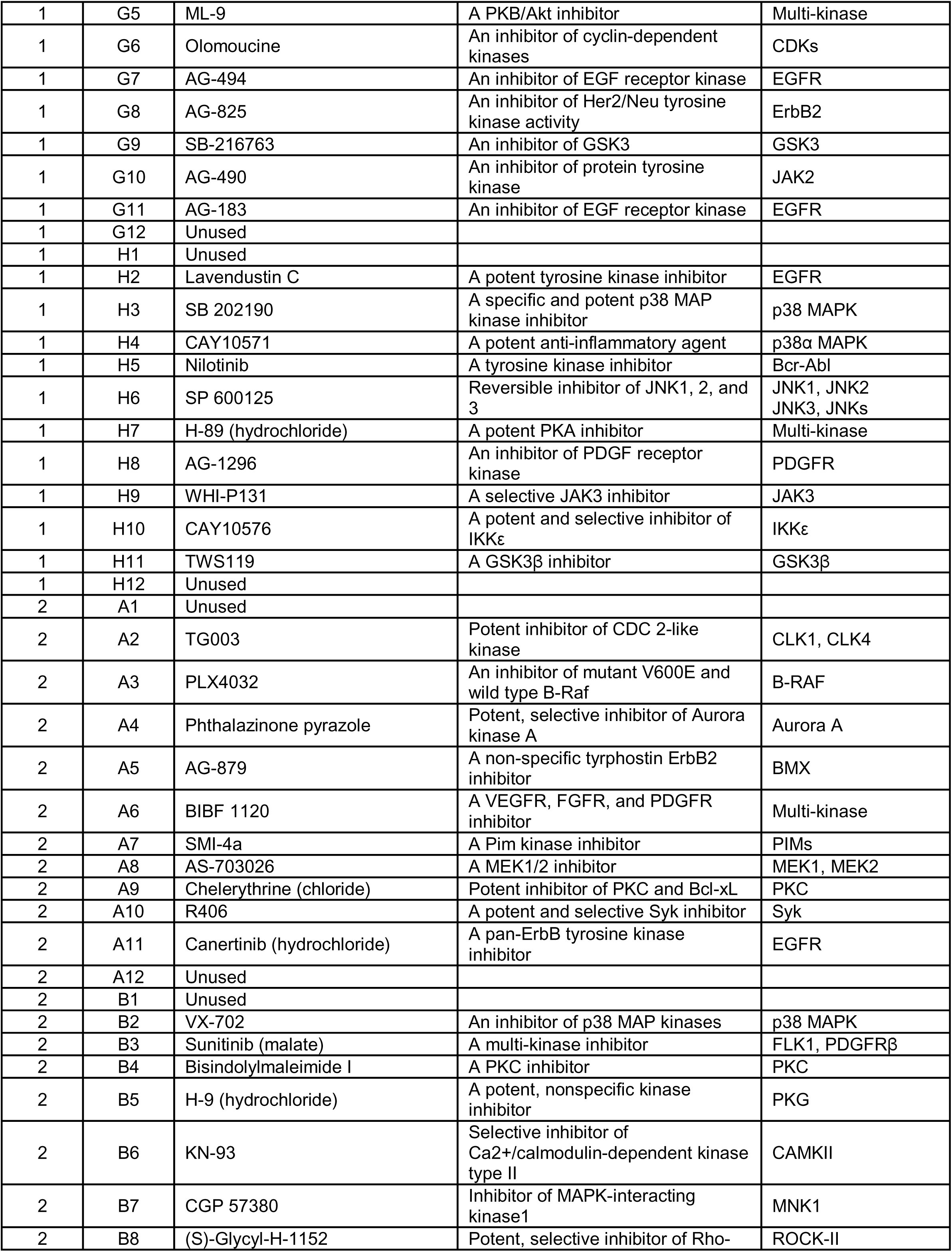
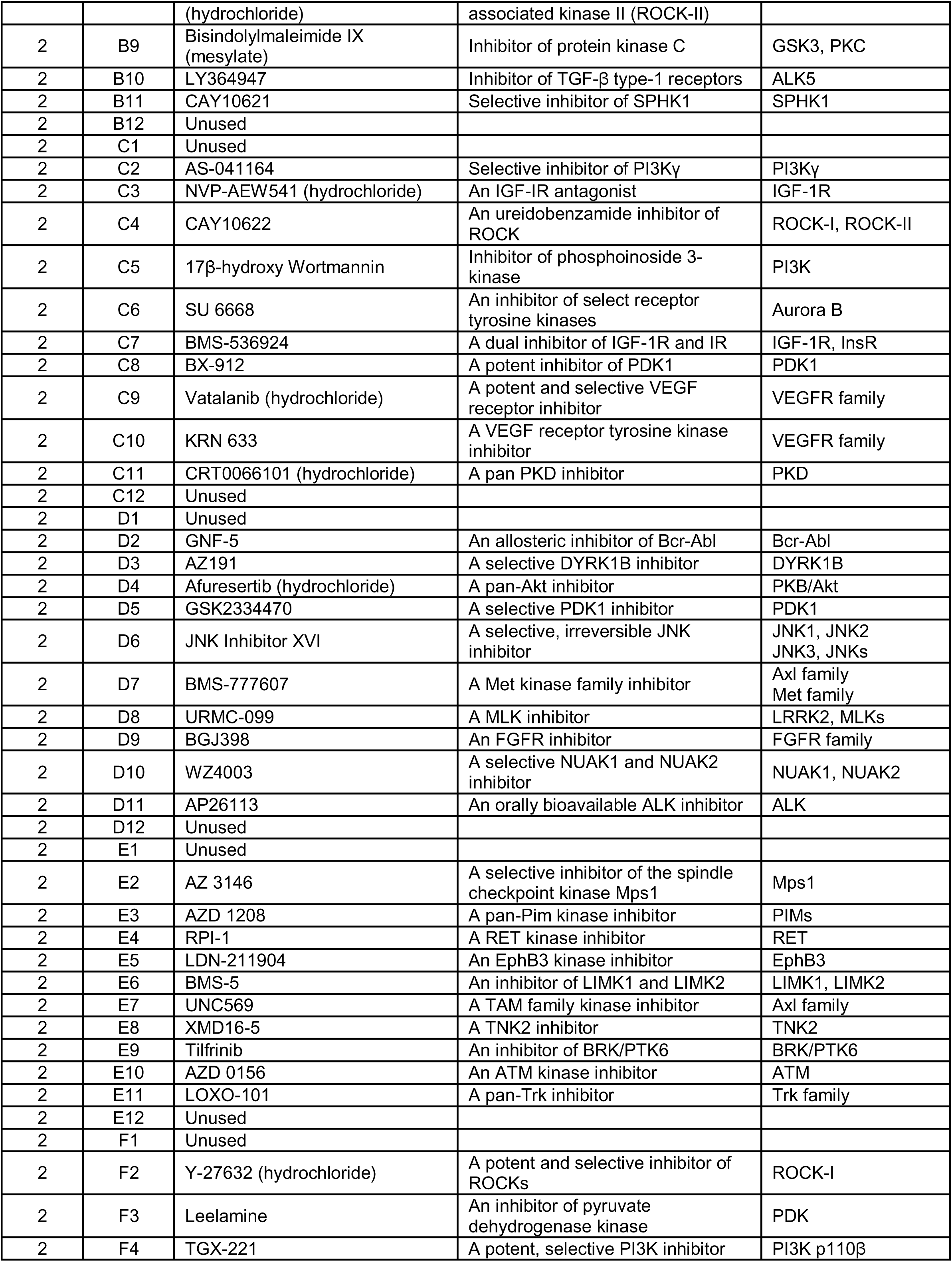
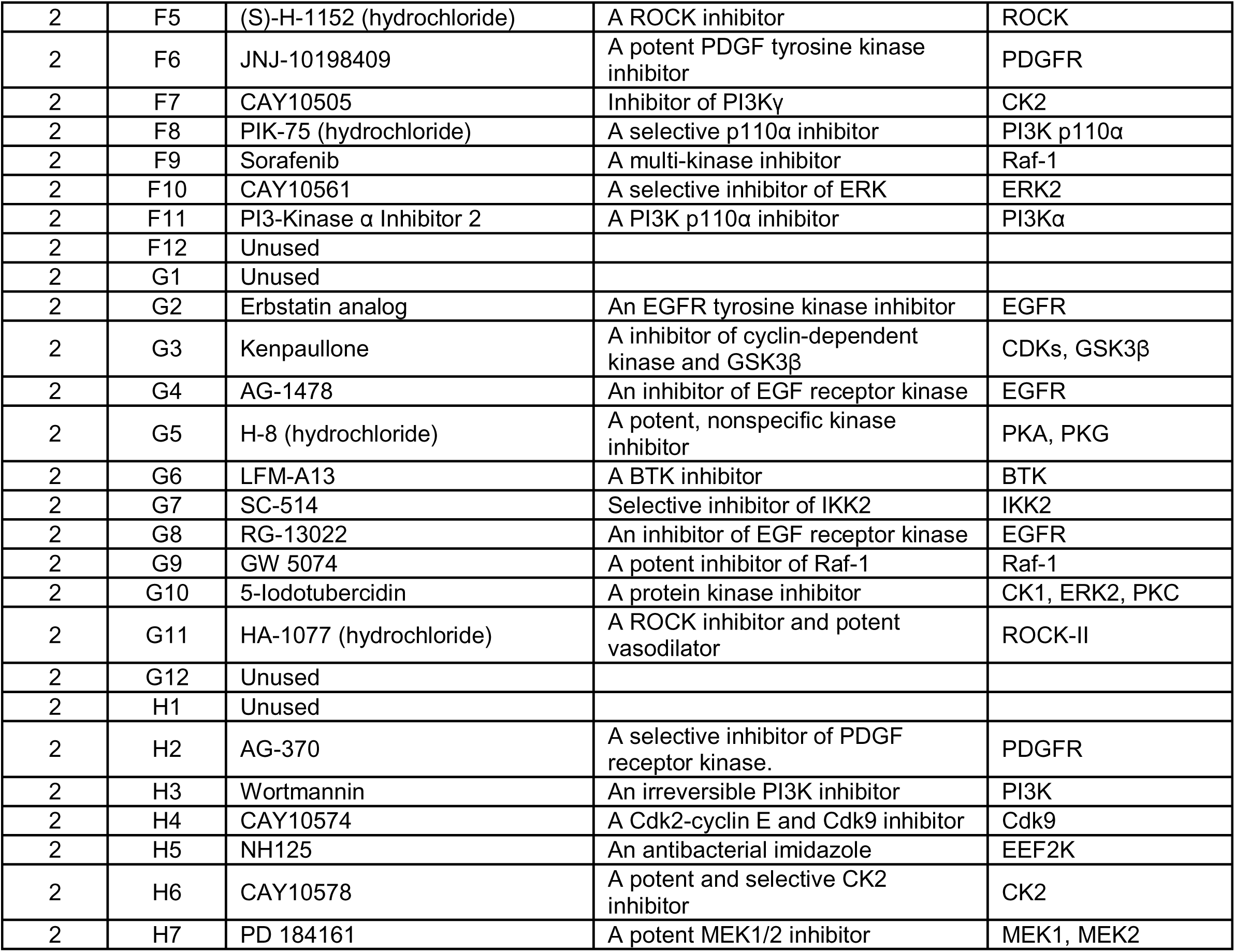
Kinase Inhibitor Library.

### Statistical Analysis

Statistical analysis was conducted using Prism 9 software (GraphPad, La Jolla, CA). The percent Tail DNA was graphed as a mean±95% confidence interval (CI). The significance was measured using an unpaired two-tailed students t-test when comparing only two groups. When comparing three or more groups, the significance was calculated by using one-way ANOVA.

## RESULTS

### Development of a 384-well CometChip

In the previous study we reported on the 96-well CometChip Platform (11) that is based on the single-cell trapping microwell approach (10). Each well can accommodate about 500 defined microwells, each containing one cell (11). With a drastic reduction in variation from sample to sample, as low as 20 comets per well are sufficient to generate robust data for a DNA damage assay when the % Tail DNA (damage) exceeds ∼15% in a 96-well CometChip assay (14). Since the microwells are evenly distributed throughout the molded agarose on a CometChip, the final throughput (the number of the total wells) of a CometChip assay is dependent on the number of wells in a microwell former (**Figure 1A**). We realized that each well of a 96-well CometChip could be split into 4 wells to generate a 384-well CometChip for higher throughput, while maintaining high sensitivity. We then developed a 384-well CometChip platform by replacing the 96-well format microwell former with a 384-well format microwell former (**Figure 1B**). The new 384-well former generated 384 individual wells on the molded agarose of each CometChip (**Figure 1B**). As a result, a 384-well CometChip increases the throughput capacity by 4-fold compared to that of a 96-well CometChip. We also designed an adapter, which can position the 384-well CometChip assembly (or the 96-well CometChip) for the use in the INTEGRA semi-automatic pipette robot for accurate transfer of compounds or cells (**Figure S1**).

### Quantifying DNA damage with the 384-well CometChip

To test if the 384-well CometChip platform can reliably detect DNA damage, TK6 cells were treated with etoposide (5 µM) or DMSO for 30 minutes in a 96-well plate at 37°C. The treated cells were then transferred into each quarter of a 384-well CometChip assembly and the CometChip assay was performed. As we predicted, the mean of the counted comets of each well of the 384-well CometChip was slightly over 100, which is about ¼ of the average of the counted comets of each well of a 96-well CometChip. There was no significant difference between the treated and untreated groups or among the four quarters (**Figure 2A**). Less than 4% (14/384) of the total wells had less than 20 comets, which is the lowest number of comets needed to generate a reliable DNA damage measurement for the CometChip assay (14). Etoposide exposure for 30 minutes resulted in an average of 32 ± 0.3% Tail DNA in the cells in each quarter of the Chip. Compared to 4.1± 0.2% Tail DNA in the DMSO control group, the 384-well CometChip assay detected a significant increase in the DNA damage from etoposide treatment (p<0.0001 for each quarter).

**Figure 2.**
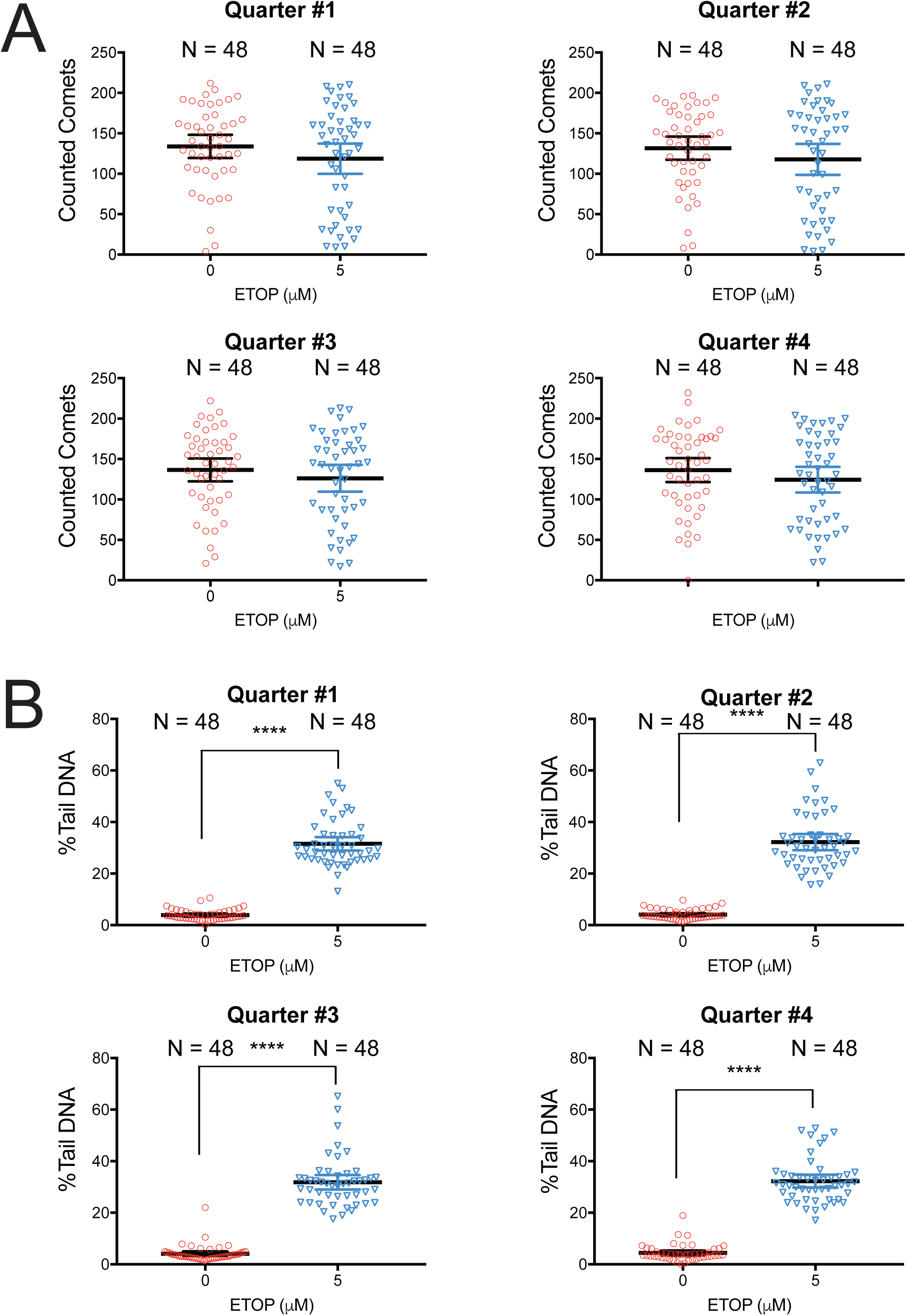
Revealing DNA damage via a 384-well CometChip. (**A**) TK6 cells were treated with etoposide (5 µM) or DMSO for 30 minutes and then transferred into each quarter of a 384-well CometChip for DNA damage analysis and to determine the number of comets of each well within each quarter of the 384-well CometChip. Each dot represents the number of the counted comets in a well of the 384-well CometChip. (**B**) The quantified DNA damage (%Tail DNA) observed after etoposide exposure in each quarter of the 384-well CometChip. Each dot represents the average DNA damage (%Tail DNA) of a well of the 384-well CometChip (****P < 0.0001)

### Sensitivity of the 384-well CometChip

Because the 384-well CometChip is designed for high throughput screening for DNA damaging events, a reasonable sensitivity is needed for the measurement of genomic insult. We compared the sensitivity of the newly designed 384-well CometChip assay with that of the 96-well CometChip assay using TK6 cells treated with a dose response of etoposide (0-10 µM) with a 2 µM ascendance. The average number of comets detected in the wells containing untreated TK6 cells in a 96 well CometChip assay was 597. Along with the increased etoposide dose, the average number of comets detected at each dose gradually decreased to 394 (**Figure 3A**). As we expected, the average number of comets detected in each well containing the untreated TK6 cells in a 384-well CometChip assay was 124, approximately ¼ of the counted comets in the 96-well CometChip assay (**Figure 3A**). The average of the total number of comets detected at each dose gradually decreased to 81, with a pattern like that of the 96-well CometChip assay (**Figure 3A**). In the 96-well CometChip assay, there was a linear relationship between the DNA damage (% Tail DNA) and etoposide dose ranging from 0 to 6 μM (R2 = 0.9825; **Figures 3 and S2**). When the cells were treated with higher doses of etoposide (8 μM and 10 μM), the DNA damage was close to the saturation level in this assay and the plot reached a plateau (**Figure 3B**). In the 384-well CometChip assay, there was also a linear relationship between the DNA damage (% Tail DNA) and the etoposide dose ranging from 0 to 6 μM (R2 = 0.993, **Figures 3, S3**) as well as a plateau for the 8 μM and 10 μM doses, like that seen in the 96-well CometChip assay (**Figure 3**). The average % Tail DNA at each etoposide dose was less than that in the 96-well assay, which may have resulted from lower levels of electric current passing through the agarose in a 384-well CometChip during electrophoresis. This change in electroosmotic flow could be a result of a tighter edge around each well, which was generated by the 384 formers. Overall, our results indicate that the 384-well CometChip assay demonstrates robust sensitivity for DNA damage analysis assays, like that seen for the 96-well CometChip assay (11).

**Figure 3.**
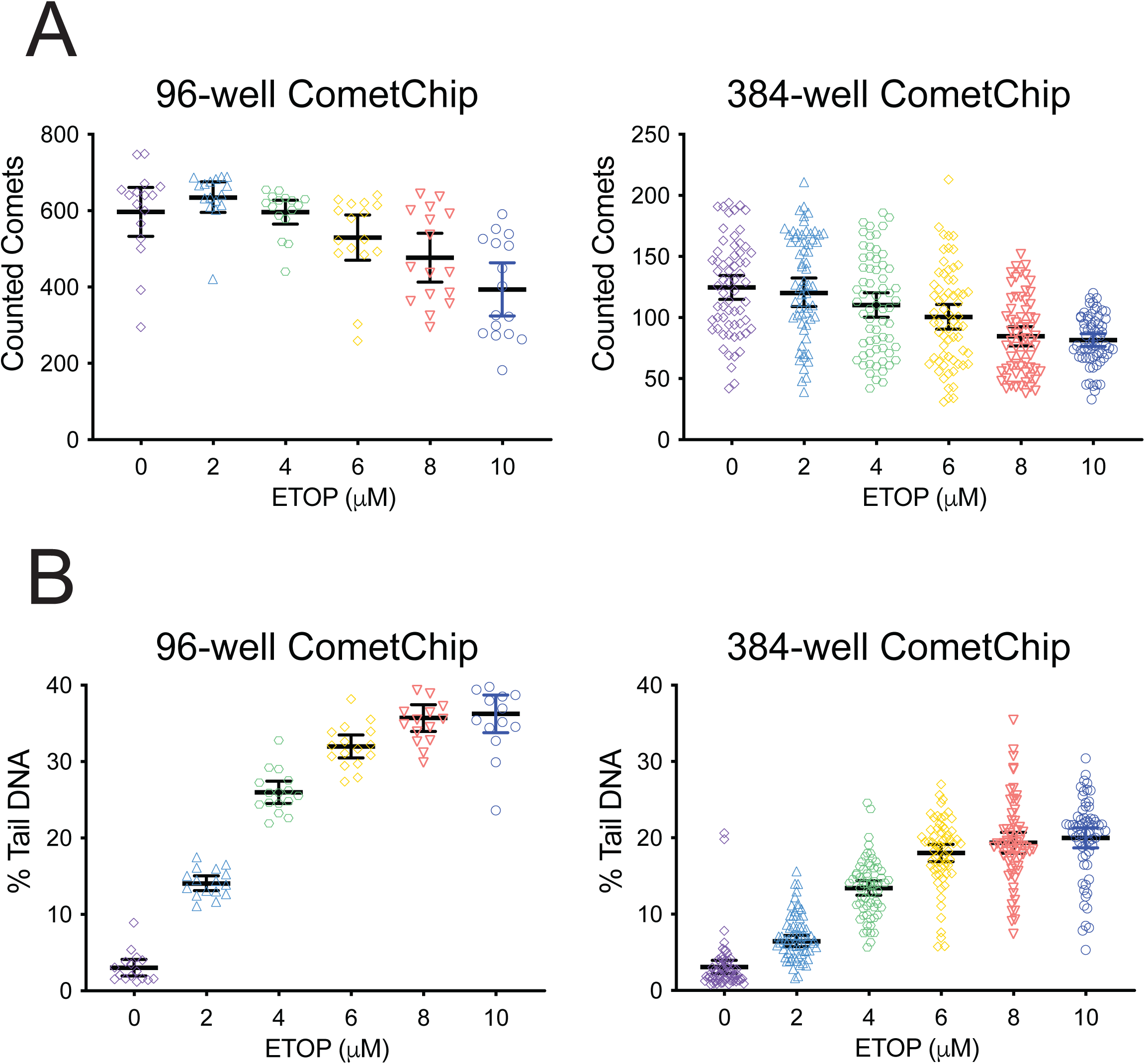
Comparison of the sensitivity of the 384-well CometChip platform to that of the 96-well CometChip platform for DNA damage evaluation. (**A**) TK6 cells were treated with etoposide (0-10 µM), with a 2 µM ascendance, for 30 minutes and the level of DNA damage was analyzed using either the 384-well platform or the 96-well platform. The number of comets within each well in a 96-well CometChip or in a 384-well CometChip is shown. Each dot represents the number of the counted comets in a well of the 96-well or 384-well CometChip. (**B**) The DNA damage (%Tail DNA) measured in cells after etoposide exposure was quantified by the 96-well CometChip assay or the 384-well CometChip assay. Each dot represents the average DNA damage level (%Tail DNA) of a well of the 96-well CometChip or the 384-well CometChip.

### 384-well CometChip assay is sensitive to loss of DNA repair capacity

It is well known that various DNA repair pathways are disrupted or deregulated in many cancers, resulting in increased mutagenesis and enhanced genomic instability, both of which promotes cancer progression (19-21). The CometChip system, therefore, is of further value to help define changes in DNA repair capacity in normal, transformed, or cancer-derived cells. We therefore investigated if the 384-well CometChip is sensitive enough to detect an alteration in DNA repair capacity when a DNA repair gene is impaired. We chose TK6 cells selectively targeted for loss (knockout) of the DNA repair scaffold gene X-ray repair cross complementing 1 (XRCC1) as a model for this proof of principle study. XRCC1 is a critical component of the base excision repair (BER) pathway and functions as a scaffold for numerous DNA repair enzymes in response to DNA single-strand breaks (22). The TK6 and TK6/XRCC1-KO cells were treated with methyl methane sulfonate (MMS), a DNA alkylating agent, at different doses for 30 minutes. Subsequently, the level of DNA damage was analyzed using the 384-well CometChip platform. Compared to the parental TK6 cells, the TK6/XRCC1-KO cells had significantly higher levels of DNA damage after MMS treatment in each quarter of the 384-well CometChip assay at each dose without an obvious difference among each quarter (**Figure 4**). At the highest dose (500 μM), MMS treatment only resulted in about 10% DNA damage (% Tail DNA) in TK6 cells. Even at the lowest dose (125 μM), MMS treatment resulted in about 30% DNA damage (% Tail DNA) in TK6/XRCC1-KO cells, which was a 3-fold increase compared to the TK6 cells treated with the highest dose (500 μM) of MMS. At this high dose (500 μM), MMS treatment induced more than 50% DNA damage (% Tail DNA) in TK6/XRCC1-KO cells, about a 5-fold increase above that of the TK6 cells (**Figure 4**). These results indicate that the loss of XRCC1, the essential scaffold protein in the BER pathway, dramatically reduces the capability of the BER pathway and demonstrates that the 384-well CometChip system was able to detect the defect in DNA repair capacity resulting from the loss of XRCC1 expression.

**Figure 4.**
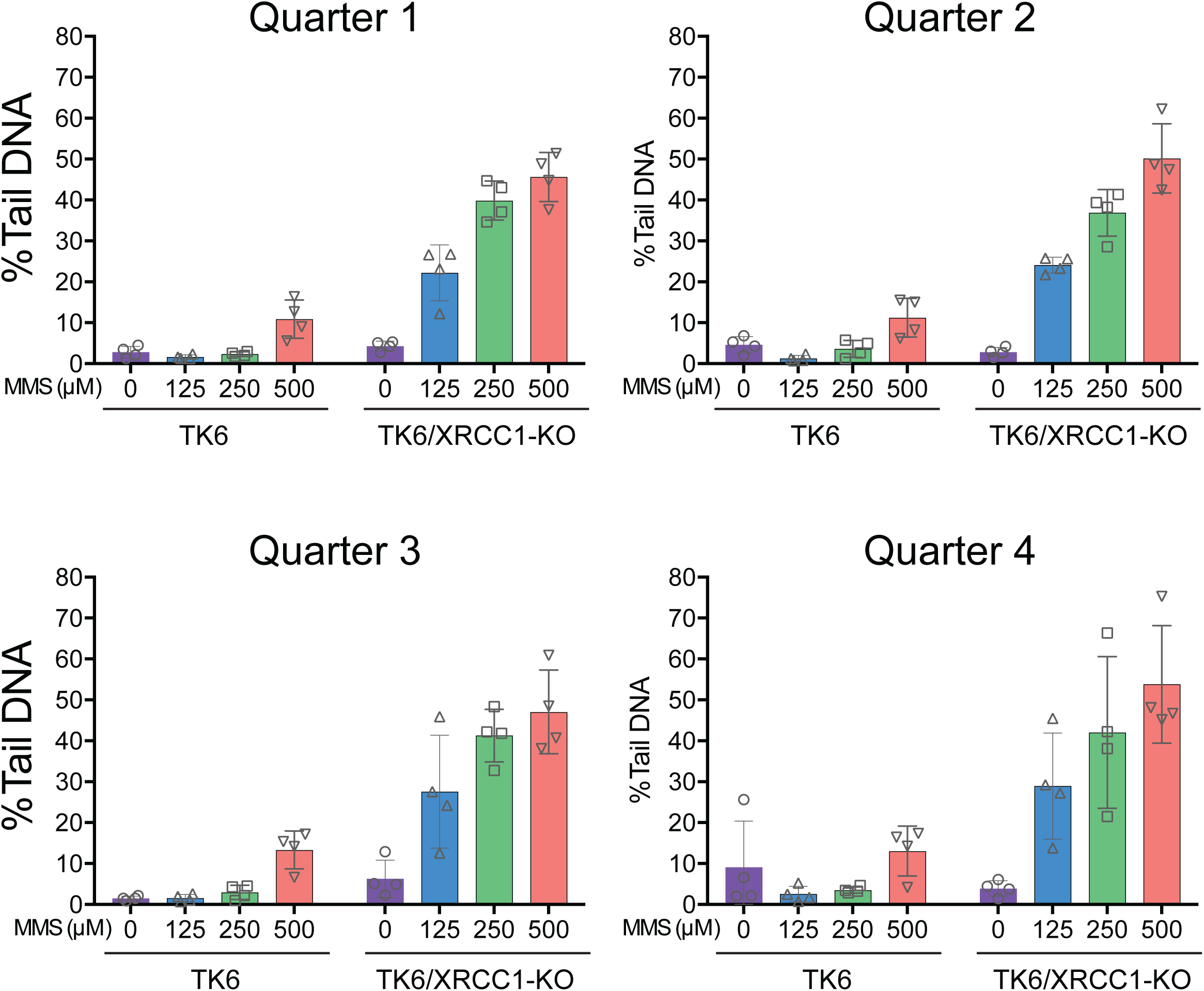
384-well CometChip assay revealed elevated DNA damage responses in cells deficient in XRCC1. TK6 or TK6/XRCC1-KO cells were treated with MMS at 0, 125, 250 or 500 µM for 30 minutes and the induced DNA damage was analyzed using a 384-well CometChip platform. The data is presented as the analysis in each quarter of a 384-well CometChip assay across 4 wells for each dose.

### 384-well CometChip assay revealed 3-methyladenine increases etoposide-induced DNA damage

Etoposide is used to treat small cell lung cancer and testicular cancer as well as other types of cancers (23). Exposure of replicating cells to etoposide generates DNA double-stand breaks by inhibiting Topoisomerase II (24). Resistance to etoposide treatment may be the result of signal transduction network activation, likely controlled by protein kinases (25-27). Therefore, we investigated the application of the 384-well CometChip platform to discover kinase inhibitors that may enhance the level of DNA damage resulting from etoposide treatment. The kinase inhibitor library we used contains 160 specific and non-specific kinase inhibitors and was chosen for this pilot experiment since it includes many compounds targeting DNA repair related signaling pathways. The library comes formatted in two 96-well plates, therefore, fitting perfectly into a 384-well CometChip assay approach. In this experiment, using the 384-well CometChip platform, 192 wells were used for the treatment with each inhibitor alone and 192 wells were combined with etoposide. Detailed information of the inhibitors is listed in the **Table 1**. We treated the TK6 cells with each kinase inhibitor (10 μM) for 30 minutes and then added etoposide (2 μM) for an additional 30-minute treatment, as compared to etoposide alone. The cells were then analyzed using the 384-well CometChip platform to score for changes in the level of induced DNA damage. As shown (**Figure 5**), overall, the cells treated with inhibitors alone (10 μM) showed no significant level of DNA damage as compared to the cells treated with DMSO as the control. In contrast, the overall level of DNA damage from the combined treatment was significantly higher than the level of DNA damage from the treatment with etoposide alone (plate #1, p = 0.0251; plate #2, p = 0.0023). The top ten hits of the combined treatment include the inhibitors targeting Yes, PIKfyve, Aurora B, PI3K, VEGFR2, PKG, SPHK1, the VEGFR family, and the EGFR pathways (**Figure 5B**). We selected the top five hits from the 384-CometChip screen and validated the result using the 96-well CometChip assay. Two out of the five hits, including Vatalanib and 3-methyladenine (3-MA), significantly increased the level of DNA damage in the cells co-treated with both 2 μM etoposide (plate #1 p = 0.008, plate #2 p = 0.006) and 4 μM etoposide (plate #1, p = 0.009, plate #2 p = 0.0009, **Figure 6A**). Furthermore, we pre-treated the TK6 cells with 3-MA (10 μM or 20 μM) for 0.5, 1 or 2 hours before etoposide (2 μM) treatment and analyzed the level of DNA damage using the 96-well CometChip assay. We found that that at each treatment condition, 3-MA significantly increased the level of cellular DNA damage following treatment with etoposide (**Figure 6B**, p < 0.0001). Altogether, our results indicate that the 384-well CometChip platform has the capacity and sensitivity for utilization in high throughput DNA damage evaluation studies.

**Figure 5.**
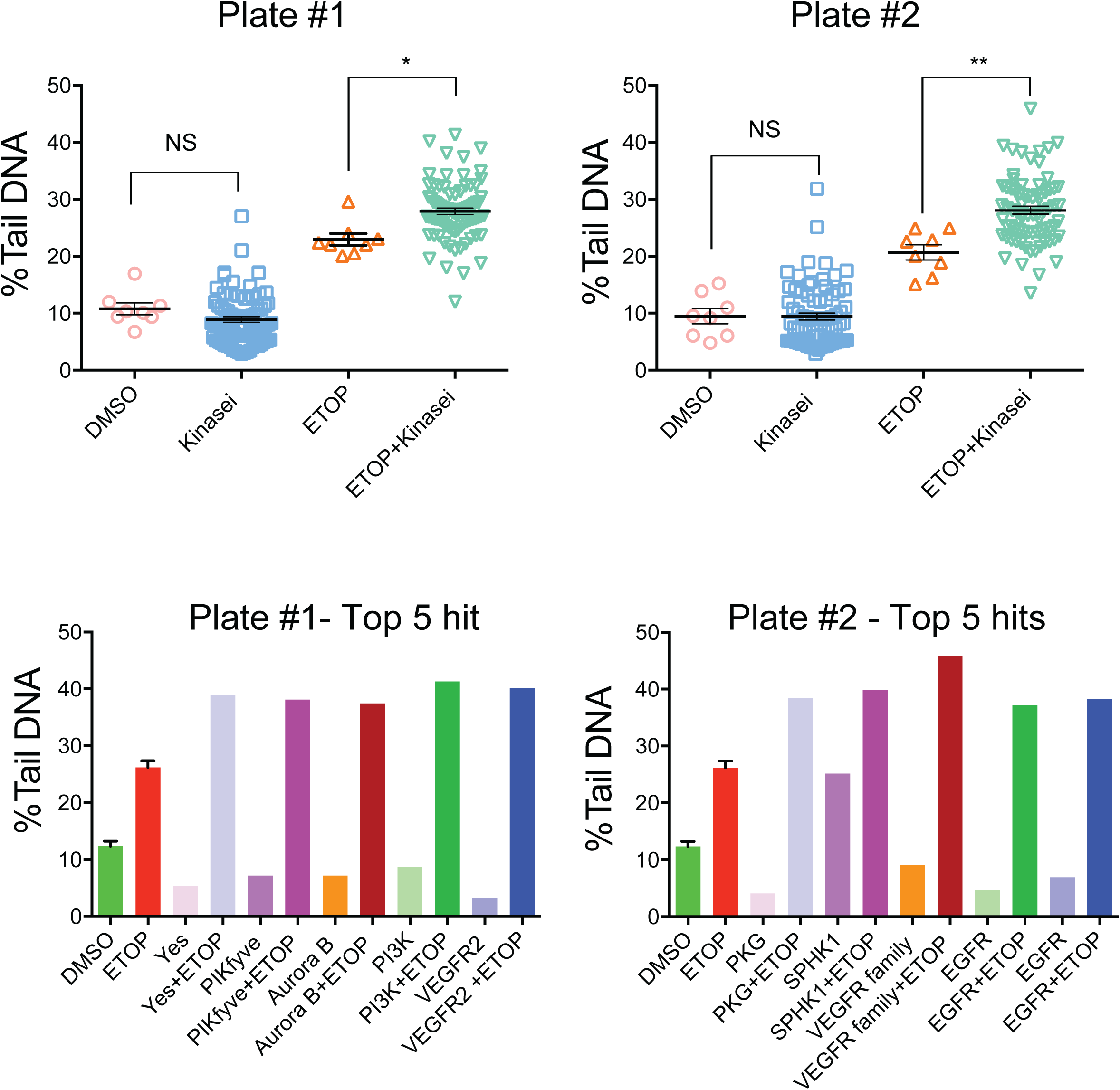
384-well CometChip assay revealed 3-methyladenine increased etoposide-induced DNA damage. (**A**) TK6 cells were pre-treated with kinase inhibitors diluted from two plates of the kinase inhibitor library for 30 minutes followed by treatment with etoposide (2 µM) for an additional 30 minutes. The DNA damage was analyzed using the 384-well platform. The overall DNA damage from DMSO, kinase inhibitor alone, etoposide alone or kinase inhibitor plus etoposide treatment for each of the kinase inhibitor sets. (**B**) The top 5 inhibitors yielding enhanced etoposide-induced DNA damage as compared to etoposide alone for each of the kinase inhibitor sets.

**Figure 6.**
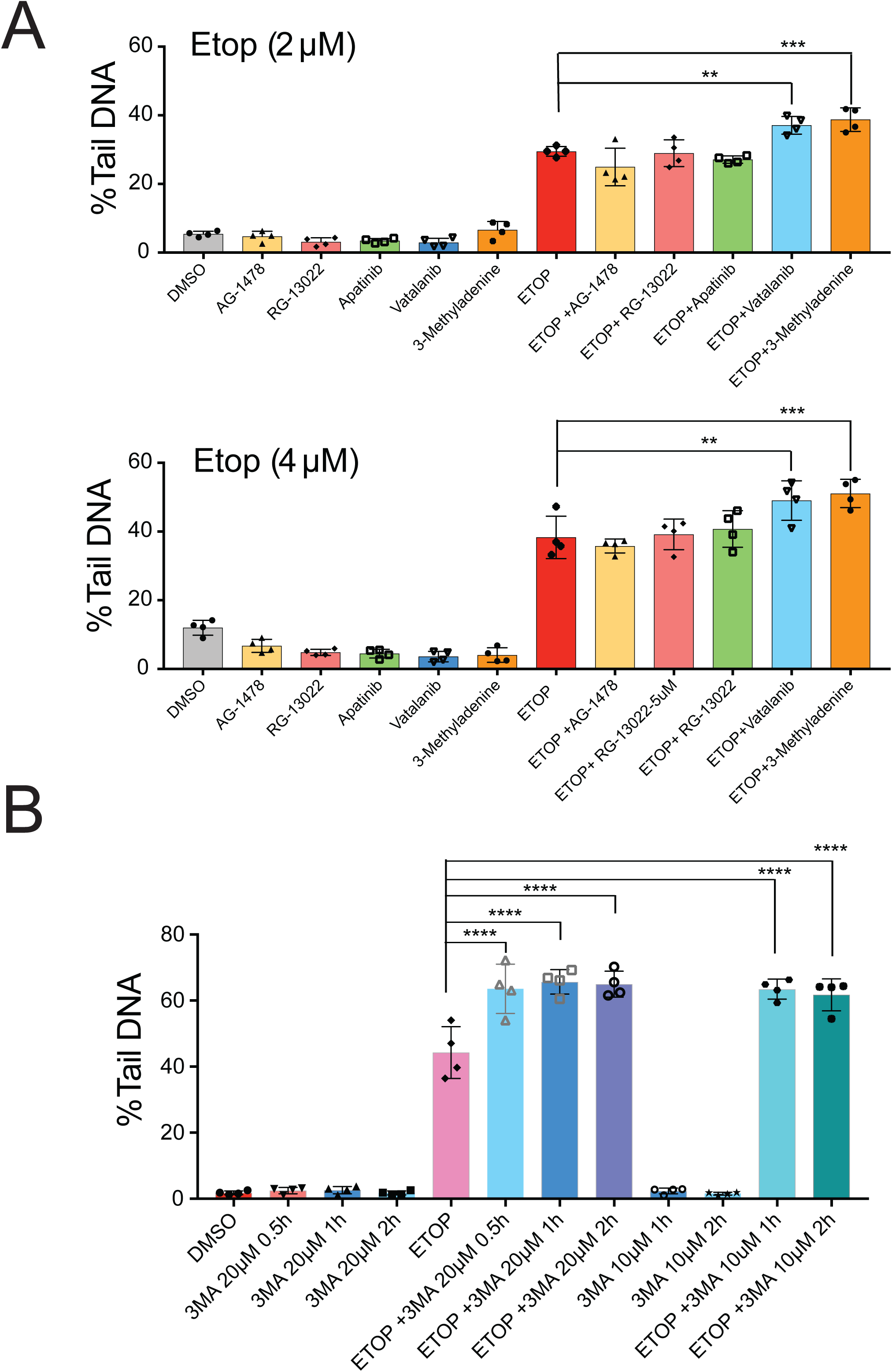
96-well CometChip assay validation of the 3-MA increase in the etoposide-induced DNA damage. (**A**) TK6 cells were pre-treated with the top 5 kinase inhibitor hits for 30 minutes followed by etoposide (2 µM or 4 µM) for an additional 30 minutes. The DNA damage was analyzed using a 96-well platform (**p < 0.01, ***p < 0.001). (**B**) TK6 cells were pre-treated with 3-MA at 10 µM or 20 µM for 1 or 2 hours followed by etoposide (2 µM) for an additional 30 minutes. The DNA damage was analyzed using a 96-well platform (****p < 0.0001).

## DISCUSSION

The comet assay is a simple and highly sensitive method for measuring global DNA damage and repair in individual cells. However, a slide-based traditional comet assay is limiting for use in large-scale experiments due to its low throughput and high data variability (28,29). Those limitations have been overcome by technical breakthroughs shown in the development of the 96-well CometChip platform (10,11), which has greatly increased the throughput and precision of the comet assay. With such an approach, a substantially lower number of comets in each well of a 96-well CometChip assay are needed to perform a reliable DNA damage analysis (14). We report here the development of a 384-well CometChip system, of which the throughput is 4-fold that of a 96-well CometChip system. Our results show a significant difference between the etoposide treated group (32% Tail DNA) and the control group (4% Tail DNA, p < 0.0001), demonstrating the ability of the 384-well CometChip to effectively measure induced DNA damage, with an average of more than 100 comets in a single well of a 384-well CometChip assay. That is nearly ¼ of the number of comets in a typical 96-well CometChip assay, which is far more than the minimum 20 comets needed for a reliable DNA damage analysis in a 96-well CometChip assay (14). Considering the increased throughput from a 384-well CometChip platform and the minimum physical distance between two comets necessary to avoid overlapping (**Figure 1**), we predict that the maximum number of total wells in a CometChip assay at the size of a universal microplate would be 1536 wells. Such an approach might yield a maximum 30 microwells per well that would surpass the minimum number of comets necessary in a CometChip assay for reliable DNA damage measurements.

The sensitivity of a 384-well CometChip assay is a critical factor for its application. Using both the 96-well CometChip and 384-well CometChip platforms, we compared DNA damage induced in TK6 cells following treatment with etoposide at a dose range of 0-10 µM. The overall dose response at the linear or saturated range related to the DNA damage (% Tail DNA) is similar between the 96-well and 384-well CometChip assays. This result indicates the sensitivity of the 384-well CometChip is consistent with that of a 96-well CometChip assay (**Figures 3B, S2**). We noticed the level of DNA damage (% Tail DNA) in a 384-well CometChip assay at each etoposide dose is smaller than that in a 96-well CometChip assay. This inconsistency is not a result of the treated cells or the performance of the experiment. After the cells were treated with etoposide in a 96-well microplate, the cells in each well were divided evenly. Half were loaded into a 384-well CometChip assembly and the remaining half in to a 96-well CometChip assembly. Therefore, the actual DNA damage of the cells treated at each dose of etoposide is the same in both assays. The subsequent procedure including lysis, electrophoresis, staining, and imaging was performed using the same solutions with the same conditions in both assays. We postulate the slightly lower level of measured DNA damage (% Tail DNA) may have resulted from the gel structure change around each well from a 384-well Comet former. We carefully measured the depth of a 96-well or 384-well CometChip former and found that the 384-well CometChip former is 1.5 mm thicker than the 96-well CometChip former. When the CometChip former was placed on the agarose gel of a CometChip by magnetic force, the 384-well CometChip former generated more pressure on the agarose gel around each well, which resulted a much thinner layer of agarose around each well. This resulted in a higher density on the gel matrix around each well. The mobility of DNA in agarose gels depends highly on the character of the gel matrix (30). An increase in agarose density results in a significant decrease in DNA migration (31). Therefore, we speculate the higher density gel matrix around each well of the 384-well CometChip changed the electroosmotic flow during electrophoresis. The cumulative effects resulted in shorter comet tails in a 384-well CometChip, showing a lower % Tail DNA after quantification. A fine modification of the voltage or an increase of gel running time during electrophoresis may be necessary for a 384-well CometChip assay to reach the same % Tail DNA as in a 96-well CometChip assay.

We additionally used the 384-well CometChip platform to highlight the critical role of XRCC1 in the BER pathway (22,32,33). Our results indicate a significant increase in the level of DNA damage when XRCC1 was depleted in TK6 cells treated with MMS. Substantial evidence indicates XRCC1 functions as a scaffold protein and physically interacts with other DNA repair factors including DNA LIGIII POLβ, APE1, PNKP, PARP1/2, OGG1 and APTX (34-39). The loss of XRCC1 fundamentally impaired the functional capacity of the BER pathway, which was demonstrated by our 384-well CometChip assay.

To demonstrate the capacity of the 384-well CometChip platform in large-scale screens, we used a small molecule protein kinase inhibitor library to investigate how kinase inhibition impacts the level of DNA damage resulting from etoposide treatment. Our results revealed that pre-treatment of TK6 cells with 3-methyladenine (3-MA) significantly increased the level of etoposide-induced DNA damage. This was further confirmed using the 96-well CometChip assay (p < 0.0001). In this study, we conducted the screen exclusively at a single dose (10 μM) and a single time point (30 min) to measure the acute DNA damage response. It is well known that the 3-MA inhibits autophagy not only by inhibiting class III PI3K but also by inhibition of class I PI3K, including those in the PI3K/AKT/GSK3β pathway (40). In most studies of autophagy inhibition, 3-MA is used at a dose of 1 to 10 mM (41-43). This is 100-fold to 1000-fold of the 3-MA dose used here (10 μM). Our findings are consistent with an earlier report that 3-MA enhanced etoposide-induced cell death of HepG2 cells by autophagy inhibition (44), although at a higher dose of 3-MA (2.5 mM), a 250-fold increase as compared to the dose used here. Therefore, we postulate that there might be additional factors, other than autophagy inhibition, that are affecting the 3-MA mediated increase in the level of DNA damage induced by etoposide. DNA lesions from etoposide treatment are mainly repaired by the non-homologous DNA end joining (NHEJ) pathway mediated by DNA-dependent protein kinase (DNA-PK) (45,46). It has been reported that AKT is involved in mediating DNA damage and repair through the NHEJ repair pathway (47-49). AKT promotes the IR-induced Ku/DNA-PK complex formation and the recruitment of DNA-PK to the DNA damage site and induces DNA-PK kinase activity and its autophosphorylation (50). Knowing 3-MA inhibits the activity of ATK (40), we speculate that the pre-treatment of a low dose of 3-MA in our assay interferes with the ATK/DNA-PK interaction axis. As a result, etoposide induced DNA double-stand breaks were less effectively repaired. Future experimental studies are necessary to validate our hypothesis.

In conclusion, we present here the first demonstration of a 384-well CometChip platform shown to be effective for large-scale DNA damage analysis studies. This is validated by using both a genetic modification as well as a chemical inhibitor approach for DNA damage evaluation. The 384-well CometChip is an effective tool for directly measuring genomic DNA damage with sensitivity comparable to that of a 96-well CometChip platform (11), but with a significant increase in the overall throughput. The capability of handling 384 samples in parallel significantly reduced the data variability and the processing time. Further, the proof-of-principle kinase inhibitor study revealed that 3-MA significantly increased etoposide induced DNA damage. Collectively, the 384-well CometChip platform has the potential to become a large-scale DNA damage analysis tool to evaluate mechanisms of DNA damage and repair as well in the development of chemotherapeutic agents that function by inducing DNA damage or inhibiting DNA repair and the DNA damage response.

## DATA AVAILABILITY

The authors declare that all data supporting the findings of this study are available within the article or from the corresponding author upon request.

## SUPPLEMENTARY DATA

Supplementary Data is available.

## ACKNOWLEDGEMENT

RWS is an Abraham A. Mitchell Distinguished Investigator. We deeply appreciate the work from our Community Advisory Board for the project, composed of the following Community Members: Latasha Farrier*, Leevones Fisher, M Ed, Barbara Hodnett**, Bobbie Holt-Ragler, DNP**, Cynthia Jackson, DNP, FNP-BC, Frewin Osteen**, Ernestine Pritchett**, and Gladys Williams* (*RA: Research Apprentice, **CHA: Community Health Advocate).

