## Supplemental Figures for "A high-throughput 384-well CometChip platform reveals a role for 3-methyladenine in the cellular response to etoposide-induced DNA damage"

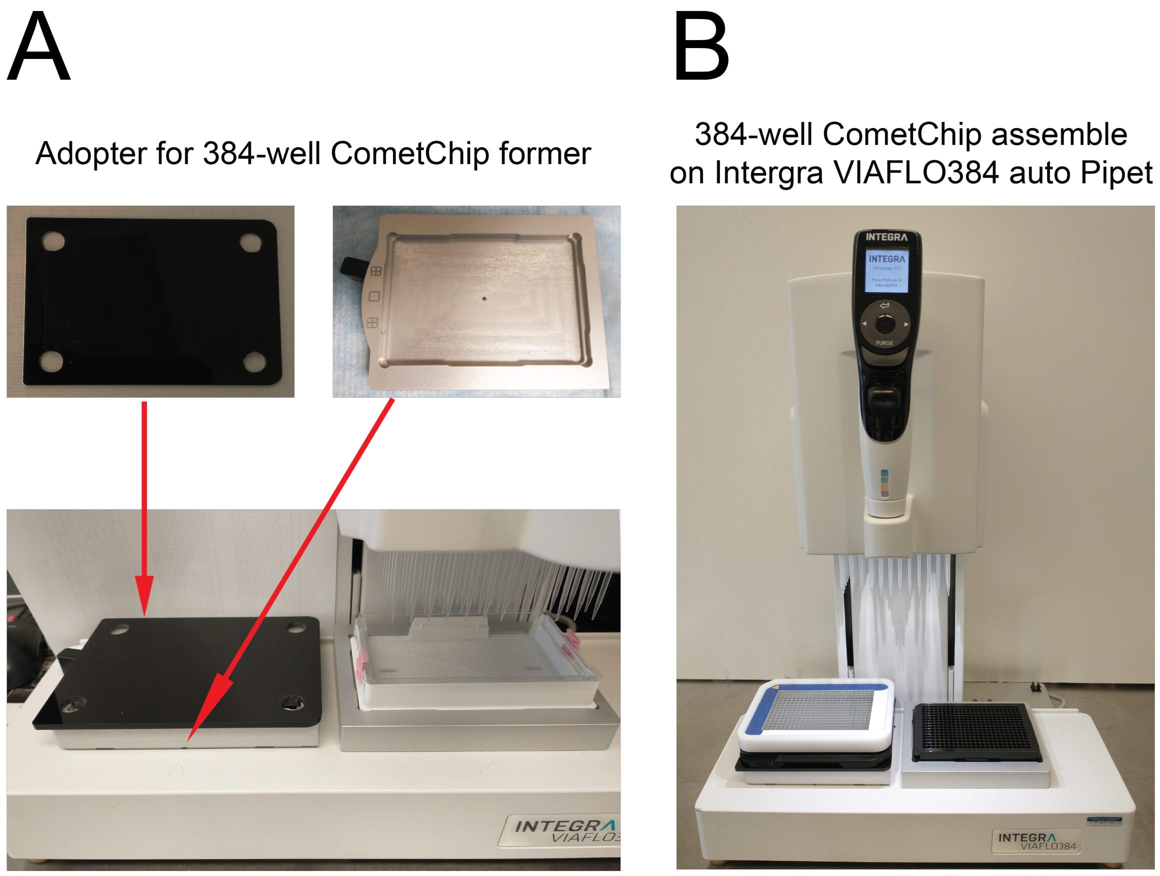


**Figure S1. Performing a 384-well CometChip assay using the Intergra Viaflo384 Auto Pipet**

**A.** An engineered adaptor to mount the 384-well CometChip assembly on the Auto Pipet system. **B**. Image showing the transfer of solutions using an Intergra VIAFLO384 auto Pipet to load the 384-well CometChip.

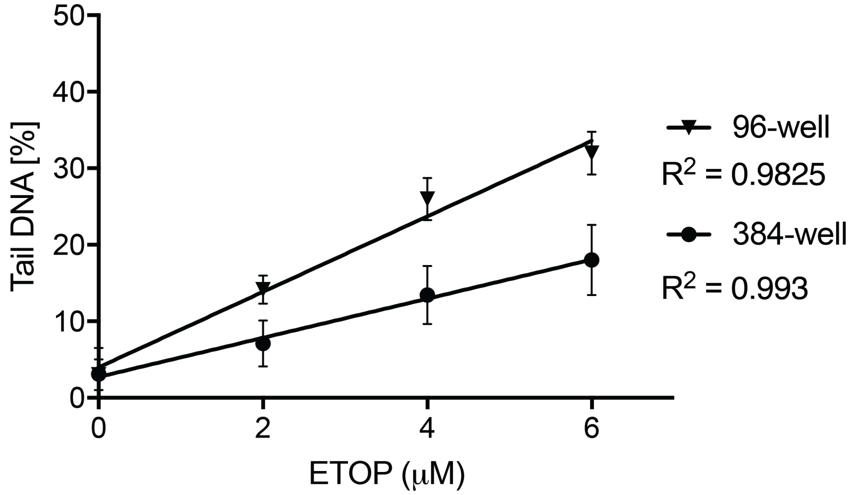


**Figure S2. Linear regression of the etoposide dose response for TK6 cells in a 96-well CometChip assay and a 384-well CometChip assay.**

The DNA damage levels in response to etoposide treatment (0, 2, 4, or, 6 µM) in the 384-well platform or the 96-well platform from **Figure 3** was re-plotted for a linear regression analysis. Line of best fit was obtained using Linear Regression analysis in GraphPad Prism. $R^2$ = 0.9825 for Etoposide doses of 0, 2, 4, and 6 µM in a 96-well CometChip. $R^2$ = 0.993 for Etoposide doses of 0, 2, 4, and 6 µM in a 384-well CometChip.
